# Integration of multidimensional splicing data and GWAS summary statistics for risk gene discovery

**DOI:** 10.1101/2021.09.13.460009

**Authors:** Ying Ji, Qiang Wei, Rui Chen, Quan Wang, Ran Tao, Bingshan Li

**Affiliations:** Department of Molecular Physiology & Biophysics, Vanderbilt University, Nashville, TN, USA; Vanderbilt Genetics Institute, Vanderbilt University Medical Center, Nashville, TN, USA; Department of Biostatistics, Vanderbilt University, Nashville, TN, USA

**Author notes:** These authors contributed equally to this work (Ran Tao, Bingshan Li).

## Abstract

A common strategy for the functional interpretation of genome-wide association study (GWAS) findings has been the integrative analysis of GWAS and expression data. Using this strategy, many association methods (e.g., PrediXcan and FUSION) have been successful in identifying trait-associated genes via mediating effects on RNA expression. However, these approaches often ignore the effects of splicing, which carries as much disease risk as expression. Compared to expression data, one challenge to detect associations using splicing data is the large multiple testing burden due to multidimensional splicing events within genes. Here, we introduce a multidimensional splicing gene (MSG) approach, which consists of two stages: 1) we use sparse canonical correlation analysis (sCCA) to construct latent canonical vectors (CVs) by identifying sparse linear combinations of genetic variants and splicing events that are maximally correlated with each other; and 2) we test for the association between the genetically regulated splicing CVs and the trait of interest using GWAS summary statistics. Simulations show that MSG has proper type I error control and substantial power gains over existing multidimensional expression analysis methods (i.e., S-MultiXcan, UTMOST, and sCCA+ACAT) under diverse scenarios. When applied to the Genotype-Tissue Expression Project data and GWAS summary statistics of 14 complex human traits, MSG identified on average 83%, 115%, and 223% more significant genes than sCCA+ACAT, S-MultiXcan, and UTMOST, respectively. We highlight MSG’s applications to Alzheimer’s disease, low-density lipoprotein cholesterol, and schizophrenia, and found that the majority of MSG-identified genes would have been missed from expression-based analyses. Our results demonstrate that aggregating splicing data through MSG can improve power in identifying gene-trait associations and help better understand the genetic risk of complex traits.

**Author summary:** While genome-wide association studies (GWAS) have successfully mapped thousands of loci associated with complex traits, it remains difficult to identify which genes they regulate and in which biological contexts. This interpretation challenge has motivated the development of computational methods to prioritize causal genes at GWAS loci. Most available methods have focused on linking risk variants with differential gene expression. However, genetic control of splicing and expression are comparable in their complex trait risk, and few studies have focused on identifying causal genes using splicing information. To study splicing mediated effects, one important statistical challenge is the large multiple testing burden generated from multidimensional splicing events. In this study, we develop a new approach, MSG, to test the mediating role of splicing variation on complex traits. We integrate multidimensional splicing data using sparse canonocial correlation analysis and then combine evidence for splicing-trait associations across features using a joint test. We show this approach has higher power to identify causal genes using splicing data than current state-of-art methods designed to model multidimensional expression data. We illustrate the benefits of our approach through extensive simulations and applications to real data sets of 14 complex traits.

## Introduction

Over the past two decades, genome-wide association studies (GWAS) have led to the discovery of many trait-associated loci. However, most loci are located in non-coding regions of the genome, whose functional relevance remains largely unclear [1]. Recent research suggested that a large portion of GWAS loci might influence complex traits through regulating gene expression levels [2, 3]. One family of methods called transcriptome-wide association studies (TWAS) has been developed to integrate GWAS and gene expression datasets to identify gene-trait associations [4]. In particular, TWAS methods like PrediXcan [5], FUSION [6], and S-PrediXcan [7] first build gene expression prediction models using reference transcriptome datasets (e.g., the Genotype-Tissue Expression (GTEx) Project [8]) and then test the associations between tissue-specific genetically predicted gene expressions and disease phenotypes using readily-available GWAS individual- or summary-level data. These methods have been widely used in practice as they facilitate the functional interpretation of existing GWAS associations and detection of novel trait-associated genes.

Gene expression is not the only mediator of genetic effects on complex traits. Splicing is of comparable importance and often functions independently of expression [3, 7, 9, 10]. The splicing process involves highly context-dependent regulation and other complex mechanisms, which could be prone to errors with potentially pathological consequences [11]. In fact, recent studies indicated that at least 20% of disease-causing mutations might affect pre-mRNA splicing [12], and splicing quantitative trait loci (sQTLs) could account for disproportionately high fractions of disease heritability [13, 14]. Despite the importance of splicing regulation, it has been understudied largely due to its complexity. Therefore, there is a pressing need to investigate trait-associated genes with effects mediated by splicing.

While gene expression can usually be summarized into one measurement per gene per tissue, there are on average eight RNA splicing events per gene per tissue [15]. To analyze splicing data, a straightforward extension of the TWAS framework for expression data is to test each genetically predicted splicing event separately and then correct for multiple testing [9, 14, 16–18]. For example, Gusev et al. [16] detected a comparable number of significant genes associated with schizophrenia from around nine times splicing events (99,562) compared to expression (10,819). While these results lend support for the importance of splicing as a genotype-phenotype link, they also suggest that there is room for appreciative power gain when information embedded in splicing events can be effectively aggregated and multiple testing burden can be dramatically alleviated.

A closely related multiple testing problem arises in TWASs when the most relevant tissue for the disease of interest is unclear, and one has to test the association between the predicted gene expression and disease outcome in each tissue separately and then apply multiple testing correction. To alleviate this multiple testing burden and improve statistical power, multi-tissue TWAS approaches like S-MultiXcan [19] and UTMOST [20] have been proposed to evaluate multiple single-tissue associations jointly by an omnibus test. Specifically, S-MultiXcan first builds gene expression prediction models in each tissue separately and then performs a chi-square test for the joint effects of expressions from different tissues on the trait of interest. To avoid collinearity issues, it applies singular value decomposition (SVD) to the covariance matrix of predicted expressions and then discards the axes of small variation. UTMOST first builds tissue-specific expression prediction models by borrowing information across tissues and then uses the generalized Berk-Jones test [21, 22] to combine associations across tissues. Recently, Feng at al. [23] proposed to use sparse canonical correlation analysis (sCCA) [24] to directly build multi-tissue gene expression features and then jointly test those sCCA features and single-tissue predicted expressions using the aggregate Cauchy association test (ACAT) [25]. They showed that this sCCA+ACAT approach could be more powerful than S-MultiXcan and UTMOST.

In this paper, we propose a multidimensional splicing gene (MSG) framework to jointly test the association between all splicing events in a gene and the trait of interest. In brief, we use sCCA to build genetically predicted multi-splicing-event features, and then perform association tests of the predicted splicing events with the trait of interest. To efficiently capture the genetic components of splicing, we use the SVD regularization approach of S-MultiXcan [19] to compute a pseudo-inverse of covariance matrix of the genetically predicted splicing events, which removes the axes of small variation. This strategy offers advantages in statistical power by reducing the degree of freedom of the chi-sqared test statistic in subsequent gene-trait association analysis.

We evaluated the performance of our MSG approach, and compared its performance with those of S-MultiXcan, UTMOST, and sCCA+ACAT through extensive simulations and real data applications. In simulations, we showed that MSG provided properly controlled type I error rates, and yielded substantial power gains over S-MultiXcan, UTMOST, and sCCA+ACAT. Real data applications using GTEx data and summary statistics from 14 complex human traits demonstrated that MSG identified on average 83%, 115%, and 223% more significant genes than sCCA+ACAT, S-MultiXcan, and UTMOST, respectively. We showcased the applications of MSG to GWAS summary statistics of Alzhimer’s disease (AD), low-density lipoprotein cholesterol (LDL-C), and schizophrenia, and found that the majority of significant splicing-trait associated genes (75%, 86%, and 89% genes for AD, LDL-C, and schizophrenia, respectively) would have been missed from expression-based analyses, highlighting the potential to incorporating splicing data into post-GWAS analyses to better our understanding of the genetic underpinnings of complex traits.

## Results

### Methods overview

Our proposed MSG method consists of two stages. In the first stage, we use sCCA to construct latent canonical vectors (CVs) by identifying sparse linear combinations of single nucleotide polymorphisms (SNPs) and splicing events that are maximally correlated with each other. In the second stage, we test for the association between each of the genetically regulated splicing CVs and the trait of interest using GWAS summary statistics. To integrate single splicing CV-trait associations into a gene-level statistic, we estimate the correlation matrix of these predicted splicing CVs using an external linkage disequilibrium (LD) reference panel. We use the SVD regularization method of [19] to determine the number of informative splicing CVs (i.e., effective degree of freedom) that explain the largest variations. Finally, we combine the associations using a chi-squared test. Fig 1 displays an overview of the MSG method (see details in the Methods section).

**Figure 1.**
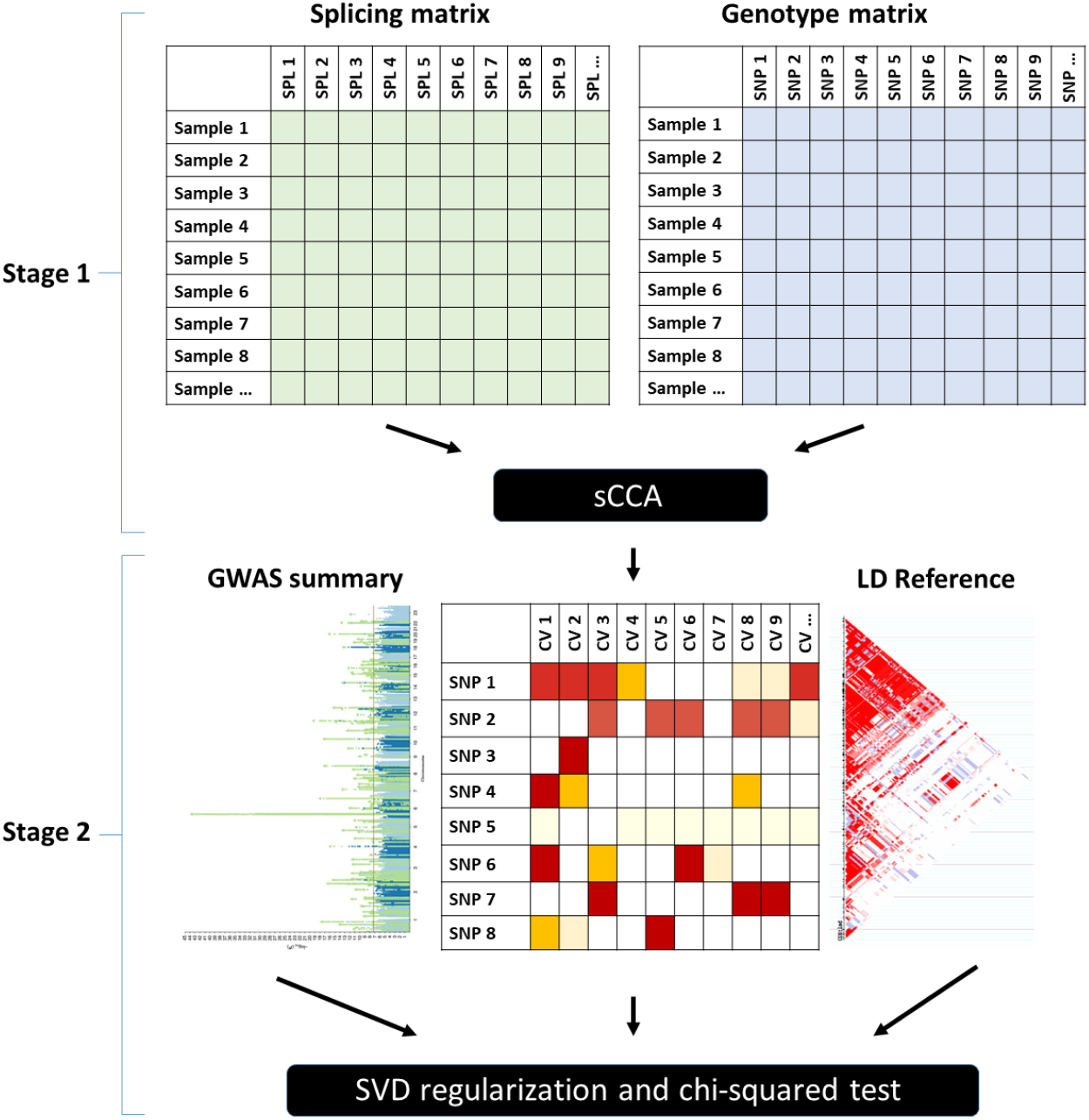
Schematic of the MSG method.

### Simulations: type I error and power analysis

We performed extensive simulations to compare the performance of MSG, S-MultiXcan, UTMOST, and sCCA+ACAT in terms of their type I error and power under various scenarios (see details in the Methods section). In the first set of simulations, we varied the number of effect-sharing splicing events (refereed to as “sharing”), the proportion of genetic variants that have non-zero effects on splicing (referred to as “sparsity”), and the cis-heritability of splicing events (referred to as 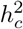). We found that MSG, S-MultiXcan, and sCCA+ACAT have properly controlled type I error rates in all scenarios, while UTMOST is slightly conservative (Table 1). Fig 2 shows that sparsity has little impact on power, yet as expected, splicing heritability increase is associated with power increase. In the second set of simulations, we defined “effect-sharing splicing events”, “non-effect-sharing splicing events”, and “trait-contributing splicing events” as splicing events that are regulated by a common set of SNPs, splicing events that are regulated by non-overlapping SNPs, and splicing events that are associated with the trait, respectively. We considered three scenarios: 1) all splicing events are trait-contributing; 2) only effect-sharing splicing events are trait-contributing; and 3) only non-effect-sharing splicing events are trait-contributing. Fig 3 shows that power increases with the number of trait-contributing splicing events for all methods, regardless of the number of effect-sharing splicing events. In both sets of simulations, we found that MSG is unanimously more powerful than S-MultiXcan, sCCA+ACAT, and UTMOST, with substantial margins.

**Table 1.**
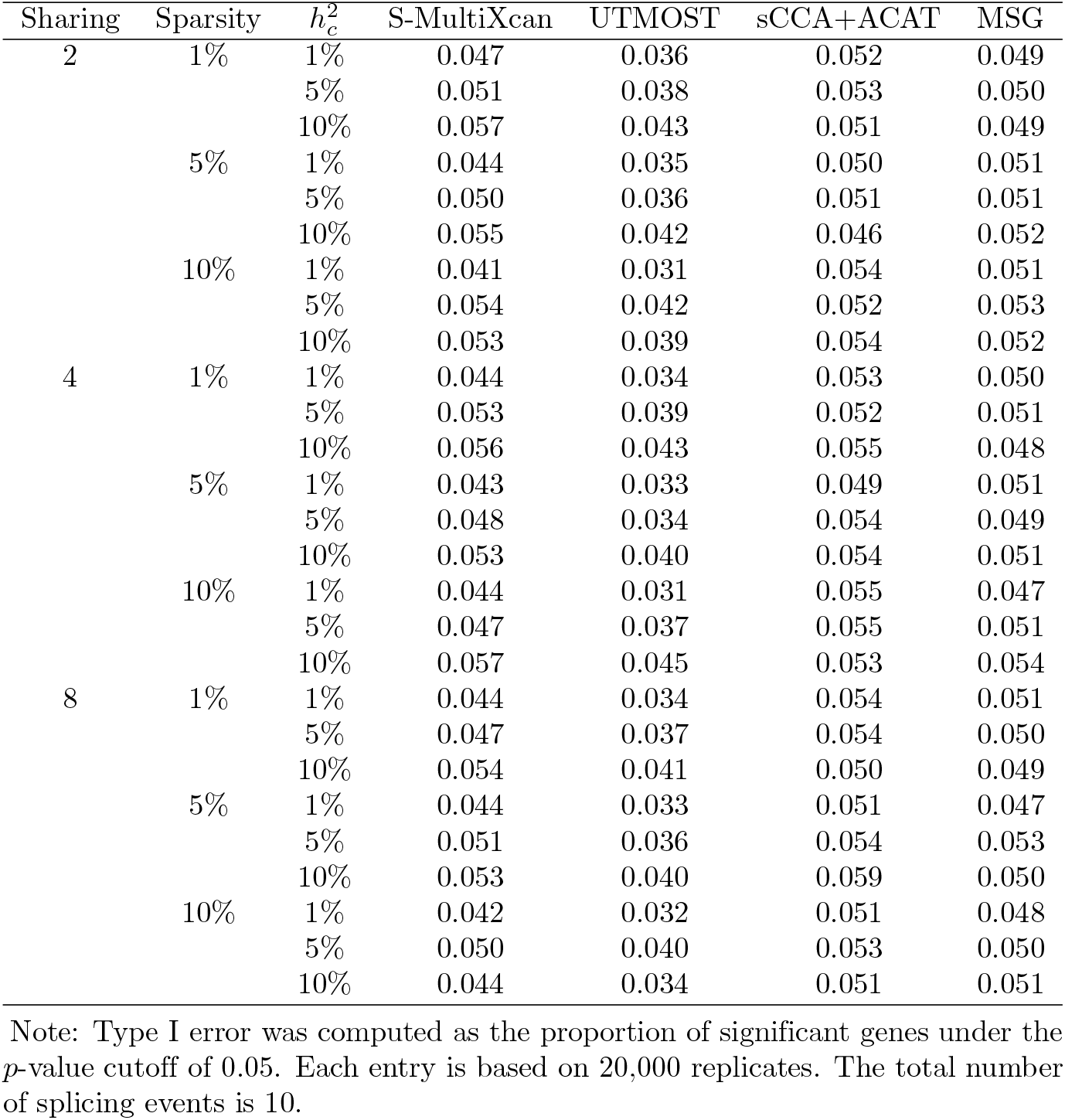
Type I error rates in the first set of simulations.

**Figure 2.**
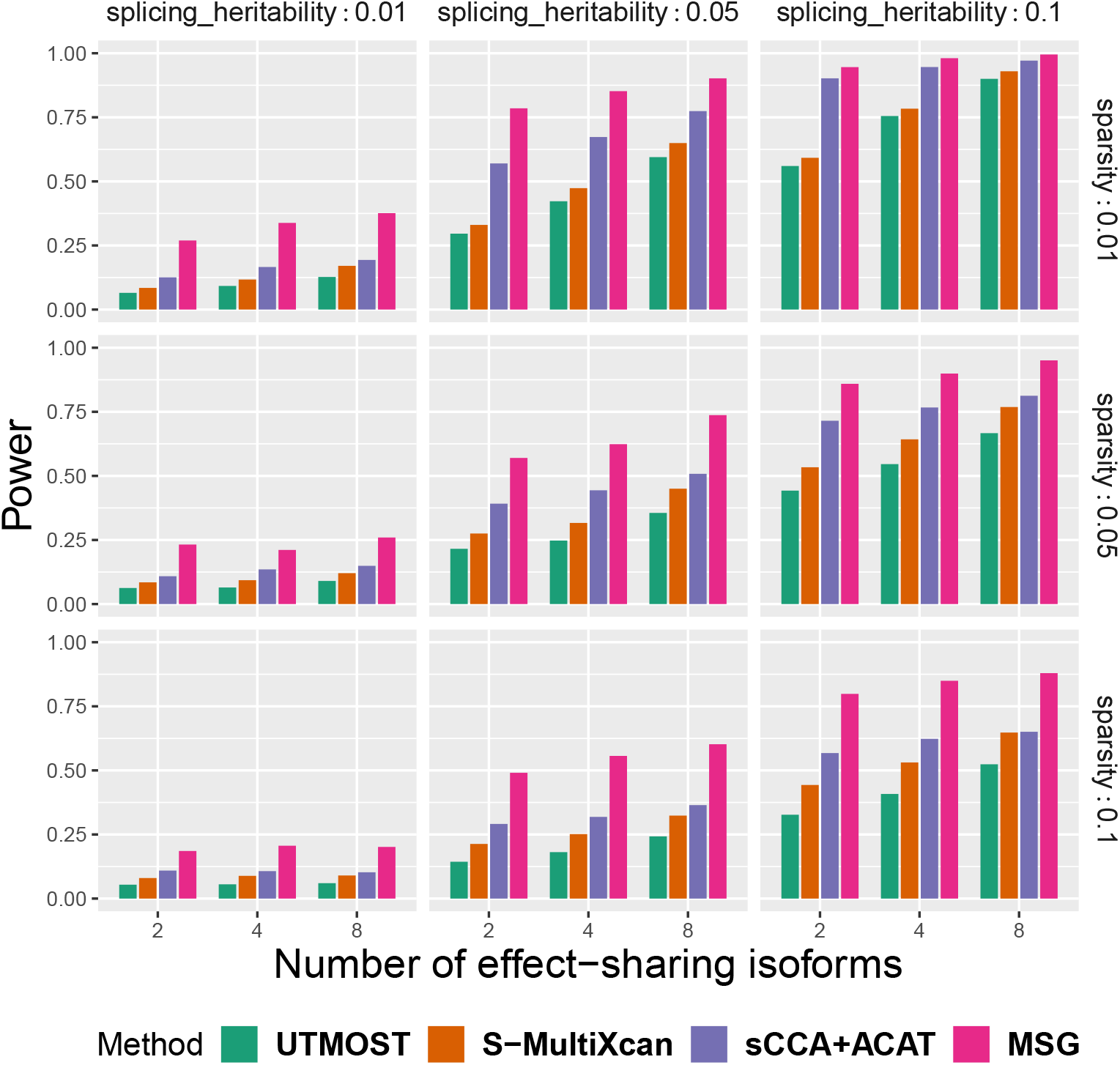
Power comparison between the S-MultiXcan, UTMOST, sCCA+ACAT, and MSG methods in the first set of simulations. With different number of effect-sharing splicing events (2, 4, 8), sparsity (0.01, 0.05, 0.1) and splicing heritability (0.01, 0.05, 0.1). The trait heritability is fixed at 0.01. For each subplot, the x-axis stands for the number of effect-sharing splicing events and the y-axis stands for the proportion of significant genes under the *p*-value cutoff of 5 × 10^−6^ across 2000 replicates.

**Figure 3.**
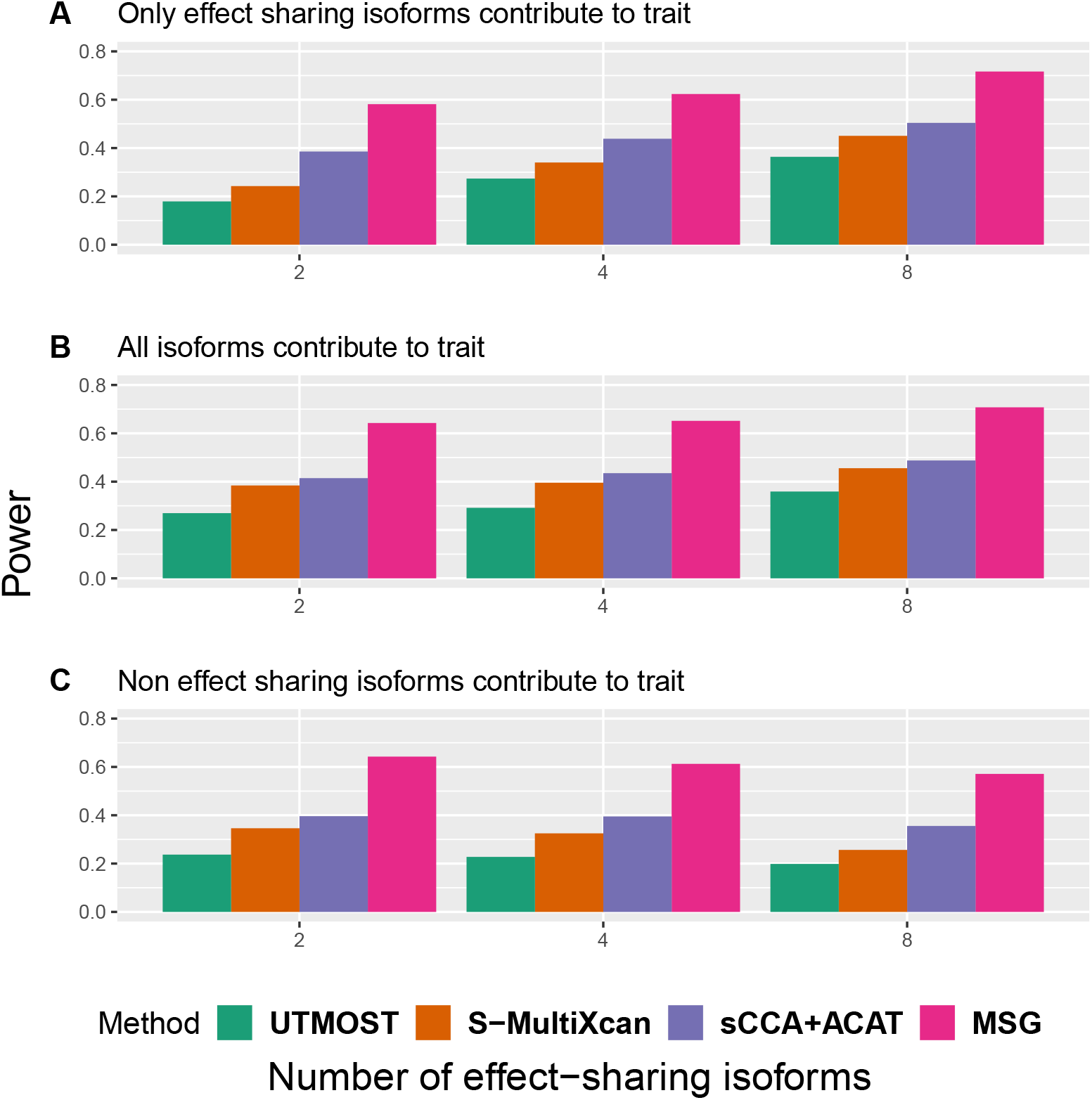
Power comparison between the S-MultiXcan, UTMOST, sCCA+ACAT, and MSG models in the second set of simulations. With different trait-contributing splicing events. For each subplot, the x-axis stands for the number of effect-sharing splicing events (2, 4, 8) and the y-axis stands for the proportion of significant genes under the *p*-value cutoff of 5 × 10^−6^ across 2000 replicates.

### Applications to complex human traits

#### Summary of applications to 14 traits

We applied MSG, S-MultiXcan, sCCA+ACAT, and UTMOST to splicing data from the GTEx project (V8 release) to obtain genetic prediction models for splicing events. We then applied the models to GWAS summary statistics of 14 complex traits to identify trait-associated genes whose genetic effects were mediated via splicing. For each trait, we chose the tissue with the top trait heritability enrichment in the respective tissue-specific annotation using linkage disequilibrium score regression [26] as previously described [20]. The sample sizes of these tissues in GTEx range from 175 (brain frontal cortex BA9) to 706 (muscle skeletal). We extracted cis-SNPs within 500 kb upstream of the transcription start site and 500 kb downstream of the transcription stop site. We selected GWASs of 14 complex traits (both quantitative and binary traits) with reasonably large sample sizes, ranging from 51,710 (bipolar disorder) to 408,953 (type 2 diabetes). When implementing S-MultiXcan, UTMOST, and sCCA+ACAT, we used the European subjects from the 1000 Genomes Project [27] as the LD reference panel to estimate the correlation matrix, following the recommendation in their original publications. When implementing MSG, we used 5,000 randomly selected European subjects from BioVU, the Vanderbilt University biorepository linked to de-identified electronic medical records [28], because MSG requires a larger LD reference panel to ensure proper type I error control (Table S1). We used Bonferroni correction to account for multiple testing across all genes for each trait separately. Table 2 shows that, at Bonferroni threshold of 0.05, MSG identified on average 83%, 115%, and 223% more significant genes than sCCA+ACAT, S-MultiXcan, and UTMOST, respectively, a substantial improvement over existing methods (see Tables S2-S16 for complete lists of significant genes identified by MSG). In particular, we examined closely the results for AD, LDL-C, and schizophrenia, with details in the next three subsections.

**Table 2.** Numbers of significant gene-trait associations across 14 human traits using S-MultiXcan, UTMOST, sCCA+ACAT, and MSG.

| Trait | Tissue | MSG | S-MultiXcan | UTMOST | sCCA+ACAT |
| --- | --- | --- | --- | --- | --- |
| AD | Brain frontal cortex BA9 | 32 | 19 | 14 | 19 |
| Bipolar disorder | Brain frontal cortex BA9 | 67 | 23 | 17 | 31 |
| Major depressive disorder | Brain frontal cortex BA9 | 23 | 5 | 3 | 0 |
| Body mass index (BMI) | Brain frontal cortex BA9 | 1757 | 786 | 497 | 704 |
| Schizophrenia | Brain frontal cortex BA9 | 458 | 203 | 145 | 234 |
| Neuroticism | Brain frontal cortex BA9 | 178 | 68 | 46 | 72 |
| Type 2 diabetes | Adipose subcutaneous | 104 | 53 | 41 | 59 |
| Total cholesterol | Liver | 202 | 109 | 66 | 118 |
| LDL-C | Liver | 200 | 108 | 69 | 120 |
| Serum urate | Liver | 87 | 63 | 50 | 64 |
| High-Density Lipoprotein Cholesterol | Adipose subcutaneous | 161 | 79 | 53 | 111 |
| Triglycerides | Adipose subcutaneous | 144 | 96 | 69 | 94 |
| Waist hip ratio adjusted for BMI | Adipose subcutaneous | 860 | 397 | 259 | 516 |
| Age at natural menopause | Muscle skeletal | 220 | 118 | 79 | 115 |

#### Application to AD

We used the brain frontal cortex BA9 splicing data from the GTEx project to build genetic prediction models for splicing events and then conducted gene-trait association analysis using the stage I GWAS summary statistics from the International Genomics of Alzheimer’s Project (IGAP) (N = 54,162) [29]. MSG, UTMOST, S-MultiXcan, and sCCA+ACAT identified 32, 14, 19, and 19 significant genes, respectively (Table 2 and Fig 4A). We observed that 26 out of the 32 MSG significant genes are within 500 kb distance to five GWAS identified lead SNPs, including the *PTK2B-CLU* locus on chromosome (CHR) 1, *SPI1* locus on CHR 11, *MS4A4A* locus on CHR 11, *PICALM* locus on CHR 11, and *APOE* locus on CHR 19 (Table S17). Among the gene-trait associations identified using MSG, 21% (7/32) were also identified by all the other three approaches; 44% (14/32) were also identified by at least one of the other approaches; and 34% (11/32) were identified by MSG only (Fig 4B). To replicate our findings, we applied these four methods to summary statistics from the GWAS by proxy (GWAX) for AD in the UK Biobank (N = 114,564) [30]. MSG, sCCA+ACAT, S-MultiXcan, and UTMOST replicated six (*MARK4, ERCC1, RELB, CLASRP, PPP1R37, CEACAM19*), two (*RELB, APOC1*), one (*RELB*), and zero significant genes, respectively, under the Bonferroni-corrected significance threshold. We compiled a list of well-known AD-associated genes (Supplementary Note Section 1A) from [31], and found that several MSG-identified AD genes are in this list (labelled in red in Fig 4C).

**Figure 4.**
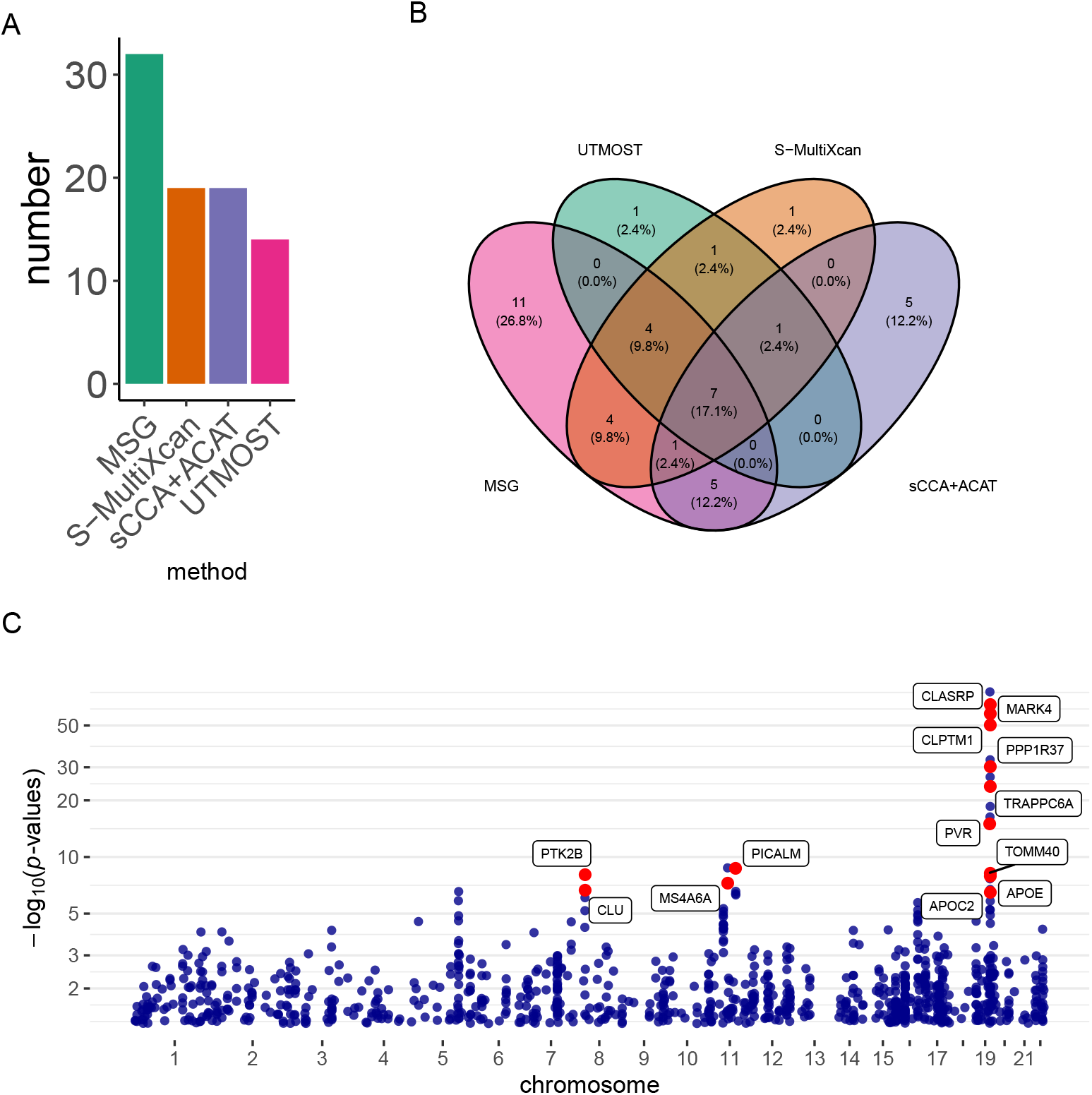
Results of the AD analysis using the IGAP stage I GWAS summary statistics. A) Bar plots of the number of significant genes using different methods; B) Venn diagram showing the overlap of significant genes identified by different methods; C) Manhattan plot for the MSG analysis. Genes with strong literature support are labeled in red.

We also conducted a conventional TWAS using S-PrediXcan with GTEx brain frontal cortex BA9 gene expression data and IGAP stage I GWAS summary statistics. We found that out of the 32 genes identified by MSG using splicing data, eight genes could also be identified by S-PrediXcan using expression data (Table S20). The remaining 24 genes would have been missed by a conventional TWAS (Fig S1). Among genes that could only be identified using splicing data, *PICALM* (MSG *p*-value = 1.93 × 10^−9^; S-PrediXcan *p*-value = 9.80 × 10^−1^) and *PTK2B* (MSG *p*-value = 7.96 × 10^−9^; S-PrediXcan *p*-value = 8.98 × 10^−1^) are two genes previously shown to be significantly differentially spliced between AD patients and healthy controls [18]. *MARK4* (MSG *p*-value = 1.31 × 10^−58^; S-PrediXcan *p*-value = 8.48 × 10^−2^) was shown to change the properties of tau [32] and has variants reported to be associated with AD and AD family history [33, 34]. Several genes in the *APOE* region are also significant in splicing but not in expression analysis: *APOE* (MSG *p*-value = 1.12 ×10^−8^; S-PrediXcan *p*-value = 2.49 × 10^−3^) is a well-known risk gene [35] for late-onset AD, with reports that alternative splicing (exclusion of exon 5) is associated with increased beta-amyloid deposition, and affects tau structure [36]; *APOC1* (MSG *p*-value = 4.65 × 10^−17^; S-PrediXcan *p*-value = 2.07 × 10^−3^) has been reported to be associated with family history of AD [37, 38]; *TOMM40* (MSG *p*-value = 5.67 × 10^−9^; S-PrediXcan *p*-value = 4.84 × 10^−1^) has been reported to have intronic variants associated with family history of AD [33] and high density lipoprotein cholesterol (HDL-C) levels [39]; *ERCC1* (MSG *p*-value = 2.32 × 10^−27^; S-PrediXcan *p*-value = 4.48 × 10^−2^), a DNA repair enzyme, has been shown to be associated with the quantification of amount of tau and implicated in AD [40].

#### Application to LDL-C

We used the liver splicing data from the GTEx project to build genetic prediction models for splicing events and then conducted gene-trait association analysis using the LDL-C GWAS summary statistics from the global lipids genetics consortium (GLGC) (N = 188,578) [41]. MSG, UTMOST, S-MultiXcan, and sCCA+ACAT identified 200, 108, 69, 120 significant genes, respectively (Table 2 and Fig 5A). We found that 102 out of the 200 MSG significant genes are within 500 kb distance to the 20 GWAS significant lead SNPs, which cluster around known SNP-level significant loci to a lesser extent than AD (Table S18). Among the gene-trait associations identified by MSG, 23% (47/200) were also identified by all the other three approaches; 37% (75/200) were also identified by at least one of the other approaches; and 39% (78/200) were identified by MSG only (Fig 5B). To replicated our findings, we applied these four approaches to summary statistics from the LDL-C UK Biobank GWAS [42] (N = 343,621) and identified 474, 223, 254, and 175 genes using MSG, S-MultiXcan, sCCA+ACAT, and UTMOST, respectively. The replication rates are high for all four methods: among the significant genes identified in the GLGC GWAS, 161 out of 200 (81%), 79 out of 108 (73%), 93 out of 120 (77%), and 52 out of 69 (75%) were replicated in the UK Biobank analysis using MSG, S-MultiXcan, sCCA+ACAT, and UTMOST, respectively, under the Bonferroni-corrected significance threshold. We compiled a list of well-known LDL-associated genes (Supplementary Note Section 1B) from [43], and found that several MSG-identified LDL-C genes are in this list, including *LPIN3, FADS3, LDLRAP1, FADS1, LDLR, FADS2* (labelled in red Fig 5C).

**Figure 5.**
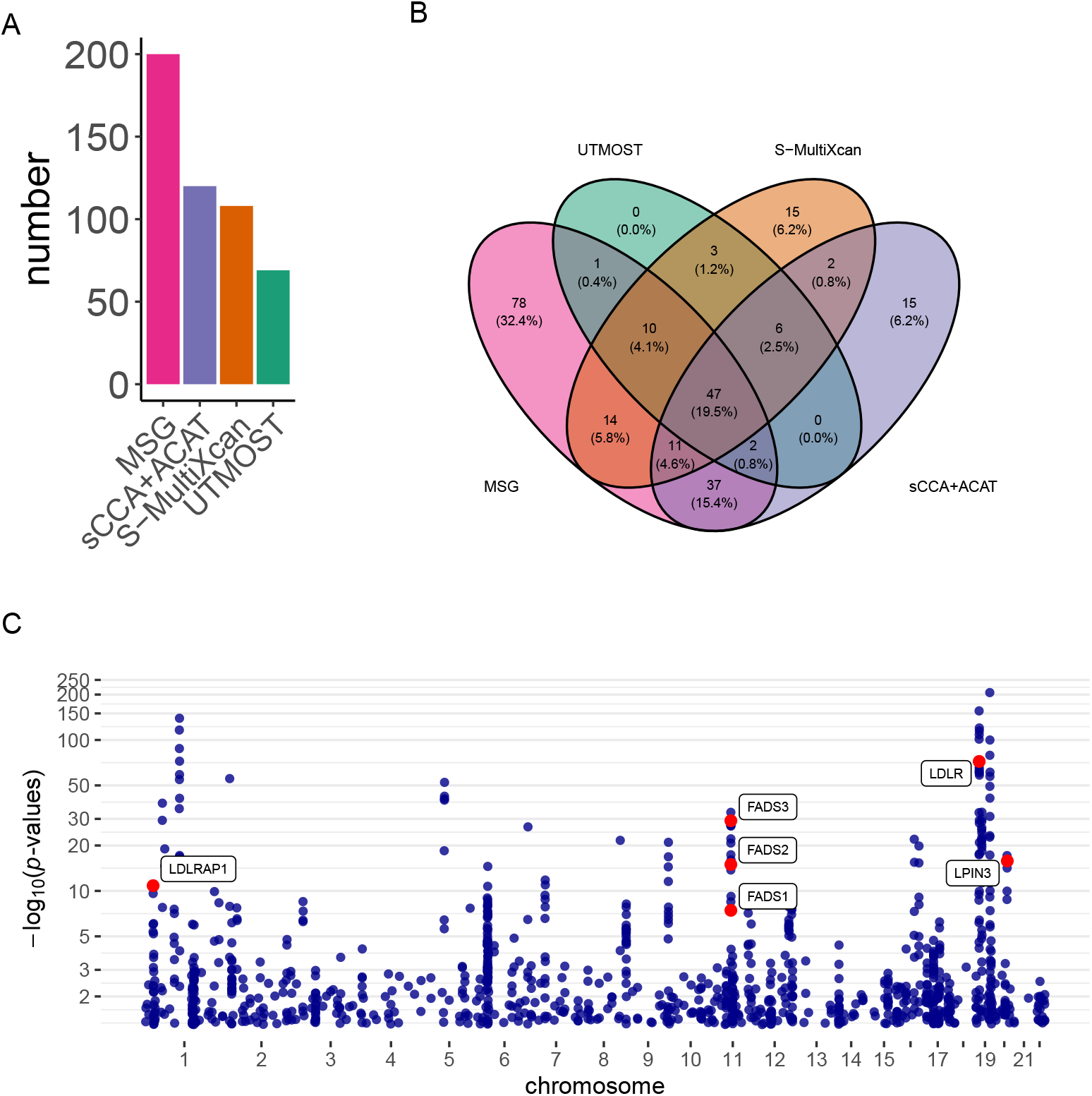
Results of the LDL-C analysis using the GLGC GWAS summary statistics. A) Bar plots of the number of significant genes using different methods; B) Venn diagram showing the overlap of significant genes identified by different methods; C) Manhattan plot for the MSG analysis. Genes with strong literature support are labeled in red.

We also conducted a conventional S-PrediXcan analysis using the GTEx liver gene expression data with the same GLGC GWAS summary statistics. We found that out of the 200 genes identified by MSG using splicing data, 27 genes could also be identified by S-PrediXcan using expression data (Table S21). The remaining 173 genes would have been missed by a conventional TWAS (Fig S2). Among genes that could only be identified via splicing: *HMGCR* (MSG *p*-value = 1.14 × 10^−40^; S-PrediXcan *p*-value = 3.00 × 10^−4^) contains variants that affect alternative splicing of exon 13 and associate with LDL-C across populations [44]; *PARP10* (MSG *p*-value = 1.51 × 10^−8^; S-PrediXcan *p*-value = 1.07 × 10^−2^) has been prioritized as a causal gene from exome-wide association analysis in more than 300,000 individuals [45]; *SMARCA4* (MSG *p*-value = 3.27 × 10^−109^; S-PrediXcan *p*-value = 6.52 × 10^−2^) has been shown to have variants associated with LDL-C levels [46], coronary heart disease susceptibility [47, 48], and myocardial infarction [49]; *LDLR* (MSG *p*-value = 4.49 × 10^−73^; S-PrediXcan *p*-value = 1.43 × 10^−1^) has been reported to be associated with statin use in the UK Biobank [50], and has intronic variants identified in familial hypercholesterolemia cases [51]; *CARM1* (MSG *p*-value = 1.05 × 10^−66^; S-PrediXcan *p*-value = 3.06 × 10^−1^) has been reported to have intronic variants associated with LDL-C and total cholesterol [52].

#### Application to schizophrenia

We used the brain frontal cortex BA9 splicing data from the GTEx project to build genetic prediction models for splicing events and then conduct gene-trait association analysis using a schizophrenia GWAS (N = 105,318) [53]. MSG, S-MultiXcan, sCCA+ACAT, and UTMOST identified 501, 222, 234, 153 significant genes, respectively (Table 2 and Fig 6A). We observe that 376 out of 501 MSG significant genes are within 500 kb distance to 76 GWAS significant SNPs (see full list of these genes in Table S19). Among the gene-trait associations identified using MSG, 18% (83/458) were also identified by all the other three approaches; 36% (165/458) were also identified by at least one of the other approaches; and 55% (253/458) were identified by MSG only (Fig 6B). Current available large-scale schizophrenia GWAS often have sample overlap, so we were unable to replicate the genes in an independent GWAS. We found that a few genes identified by MSG had been reported to influence schizophrenia risk via splicing (Supplementary Note Section 1C), including *SNX19* [54], *AS3MT* [55], and *CYP2D6* [54] (Fig 6C).

**Figure 6.**
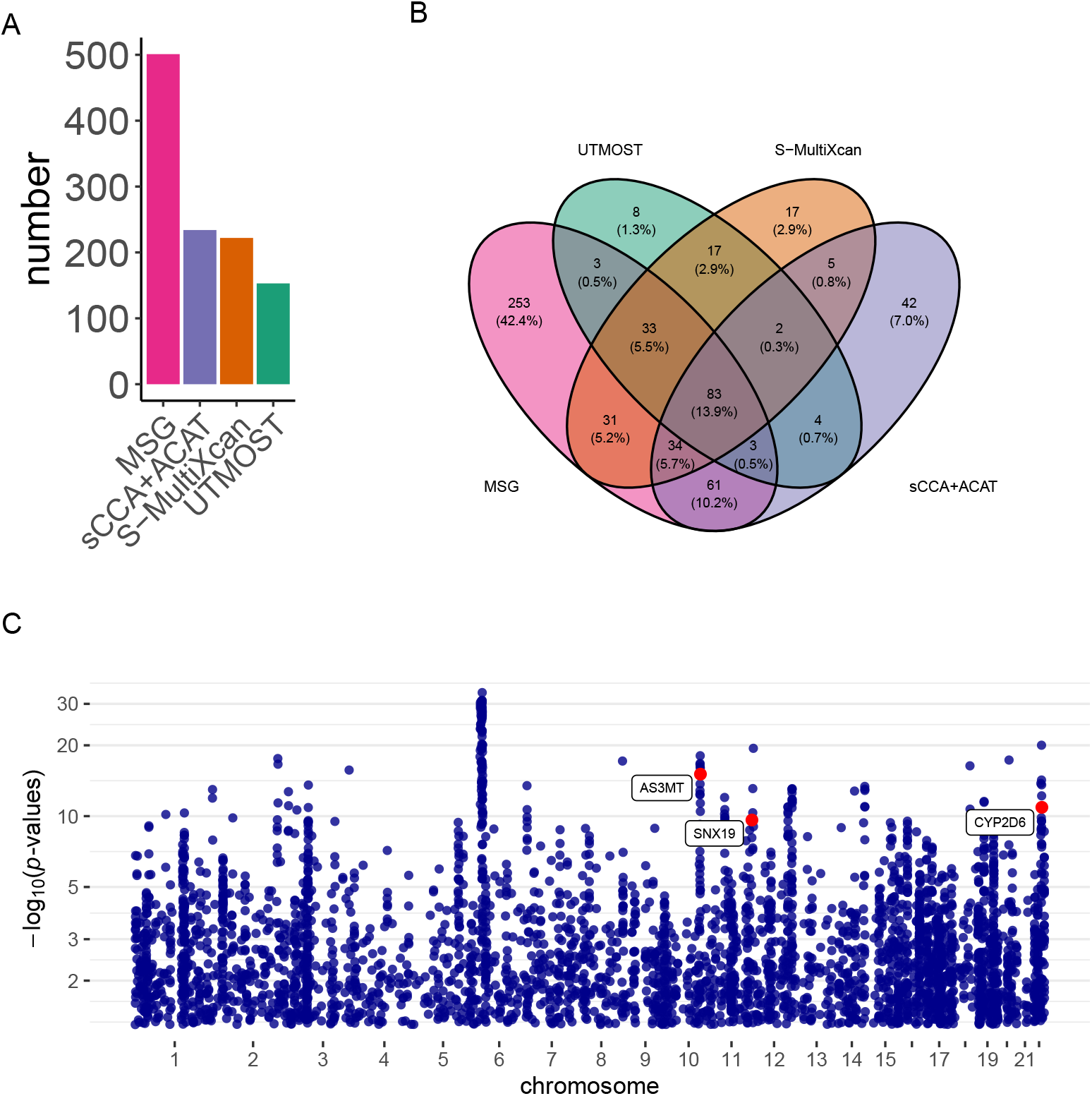
Results of schizophrenia analysis. A) Bar plots of the number of significant genes using different methods; B) Venn diagram showing the overlap of significant genes identified by different methods; C) Manhattan plot for the MSG analysis. Genes with strong literature support are labeled in red.

We also conducted a conventional TWAS using S-PrediXcan, GTEx brain frontal cortex gene BA9 expression data, and the same GWAS summary statistics. We found that out of the 458 genes identified by MSG using splicing data, 55 genes could also be identified by S-PrediXcan using expression data. Due to the complex haplotype and LD structure of the major histocompatibility complex (MHC) region, we summarized the results for genes in and outside of the MHC region separately. In the MHC region, 30 genes overlapped between 33 genes identified by S-PrediXcan and 101 genes identified by MSG (Table S22). Genes with literature support to be associated with schizophrenia that could only be identified using splicing data in the MHC region includes *NOTCH4* (MSG *p*-value = 8.35 × 10^−29^; S-PrediXcan *p*-value = 8.19 × 10^−2^) [56], *TRIM26* (MSG *p*-value = 4.64 × 10^−14^; S-PrediXcan *p*-value = 4.40 × 10^−1^) [57], and *ZSCAN9* (MSG *p*-value = 4.64 × 10^−14^; S-PrediXcan *p*-value = 4.40 × 10^−1^) [58]. Outside of the MHC region, 25 genes overlapped between 58 genes identified by S-PrediXcan and 400 genes identified by MSG (Table S22). Among genes that could only be identified using splicing data (Fig S3), *SNX19* (MSG *p*-value=2.27 × 10^−10^; S-PrediXcan *p*-value=2.38 × 10^−3^) has been reported to have schizophrenia risk-associated transcripts, defined by an exon-exon splice junction between exons 8 and 10 (junc8.10), which is predicted to encode proteins lacking the characteristic nexin C terminal domain [59]; *GRIA1* (MSG *p*-value=1.28 × 10^−8^; S-PrediXcan *p*-value=1.62 × 10^−5^) has been reported to be a schizophrenia risk gene [60]; *CACNA1C* (MSG *p*-value=9.35 × 10^−10^; S-PrediXcan *p*-value=5.44 × 10^−1^) and *CACNA1G* (MSG *p*-value=2.45 × 10^−6^; S-PrediXcan *p*-value=7.15 × 10^−1^) encode calcium voltage-gated channel subunit and have been implicated in multiple studies to be a risk gene associated with schizophrenia [60, 61]; and *PPP1R16B* (MSG *p*-value=4.86 × 10^−18^; S-PrediXcan *p*-value=8.62 × 10^−1^) has been reported to be associated with schizophrenia in several populations [60, 62] and multiple psychiatric disorders [63, 64].

## Discussion

While there is extensive research on trait-associated gene discovery based on gene expression using methods like S-Predixcan and FUSION and their multidimensional variants like S-MultiXcan, UTMOST, and sCCA+ACAT recently, there has been few studies on trait-associated gene discovery using splicing data so far. Splicing data present unique challenges due to its multidimensional nature, which demands the development of efficient analytic approaches. In this paper, we proposed MSG, a framework to construct cross-splicing event models using sCCA to boost power in identifying genes influencing traits via splicing. Through simulations, we showed that MSG has proper type I error control and superior power compared to current state-of-the-art approaches, e.g. S-MultiXcan, UTMOST, and sCCA+ACAT. In real data applications, MSG identified on average 83%, 115%, and 223% more significant genes than sCCA+ACAT, S-MultiXcan, and UTMOST, respectively, across 14 complex traits. We highlighted our findings on AD, LDL-C, and schizophrenia, and found independent literature support for MSG-identified genes, showcasing MSG’s advantage of capturing novel risk genes mediated via splicing.

Through MSG, we found a considerable number of trait-associated genes that were not identified from S-PrediXcan using expression data, demonstrating the complementary roles of genetic regulation through splicing and expression on trait variation and disease susceptibility. The number of genes identified by MSG using splicing data is usually larger than that identified by S-PrediXcan using expression data. A few factors may contribute to this phenomenon. One factor is that splicing is highly prevalent, affecting over 95% of human genes [12]. It provides the possibility of cell type- and tissue-specific protein isoforms, and the possibility of regulating the production of different proteins through specific signaling pathways [65]. Another factor is that the rich multidimensional splicing information may yield higher power to detect gene-trait associations compared to one-dimensional expression information. It was shown that the power of conventional TWASs increase to a maximum when the sample size of the reference transcriptome dataset exceeds 1000 [6]. As most tissues in GTEx have sample sizes less than 1000, the sample size of a target tissue may be too small to yield enough power for expression data-based TWAS analysis, but may be sufficient to detect associations for multidimensional splicing data analysis. Thus, we believe that splicing data may offer unique opportunities to study genetic risk of complex traits, and view our method as an important step toward using sQTLs for GWAS interpretation and gene discovery.

We observed 83%–223% increase in the number of trait-associated splicing genes identified by MSG compared to established methods like sCCA+ACAT, S-MultiXcan, and UTMOST. The relative increase of power using MSG can be attributed to several factors. Specifically, the MSG models tend to be less sparse (i.e., include more SNPs with non-zero weights) than the S-MultiXcan and UTMOST models and explain more variability in splicing variation. As a result, we found that in MSG, more genes are “testable” than S-MultiXcan and UTMOST. For example, there are 1041 genes not testable by S-MultiXcan or UTMOST but testable by MSG using the brain frontal cortex BA9 splicing data from the GTEX project. MSG is also substantially more powerful than sCCA+ACAT, despite the fact that both use SCCA to build genetically regulated splicing models. We speculate that it may be due to the following reasons: 1) MSG directly uses the sCCA-generated CVs for association tests, while sCCA+ACAT retrains the splicing CV models using elastic net, which tends to generate models that are more sparse and captures less splicing variation; 2) MSG chooses the number of CVs to be included in the association test in an adaptive manner using the SVD regularization approach of [19], while sCCA+ACAT uses three CVs throughout, which may not be optimal for all genes and tissues; 3) MSG fully incorporates information from multiple CVs using a multi-degree-of-freedom chi-square test, while the ACAT test directly combines p-values and thus could entail information loss.

Because the MSG models tend to be less sparse compared to alternative methods, they require a larger reference panel than the commonly used 1000 Genomes European samples to ensure accurate LD calculation and proper type I error control. We conducted simulation studies using LD reference panels of different sizes when performing gene-trait association analysis using the summary statistics of a GWAS with 50,000 samples and found that a reference panel of 5,000 individuals is adequate for MSG (Table S1). To construct this large reference sample in practice, we randomly selected 5,000 samples of European descent in BioVU [28]. When such LD reference panels are not available, one may still use the 1000 Genomes samples for initial screening purposes, but more stringent validation will be needed to follow up with the candidate genes identified.

There are several limitations in our study. First, we focused on single-gene, single-trait analyses of splicing data, and there are exciting opportunities for methods development and gene discovery in multi-tissue, multi-trait, multi-gene, and cis and trans effects analyses [23, 66, 67]. Second, we used the GTEx transcriptome data from adult bulk tissues. Consequently, findings driven by differences in cellular composition or developmental stages cannot be fully resolved. As splicing is likely to be tightly regulated, the association of splicing implicated genes with traits in different cell types or developmental stages remains to be studied. Third, like other TWAS-type approaches, results from our method need to be interpreted with caution: they do not implicate causality. Further causal analysis using methods like FOCUS [68] and experimental validation are needed to determine causal genes.

## Conclusion

By integrating multidimensional splicing information with GWAS summary statistics, we are able to pinpoint candidate risk genes associated with common traits via splicing. This approach can potentially be extended to integrate molecular data beyond splicing, such as epigenetic data. With the increasing availability of GWAS summary statistics of many complex traits and molecular data, we believe that our framework and its extensions will enable us to better understand how genes influence complex traits through diverse regulatory effects.

## Methods

### MSG framework

In this study, we use splicing and genotype data from the GTEx project and GWAS summary statistics of the traits of interest to identify splicing-trait-associated genes. For a given gene, let *n, p*, and *q* denote the sample size, number of SNPs in the cis-region of the gene (i.e., a 1-Mb window around the transcription start sites of a gene), and the number of splicing events, respectively, in GTEx. We note that *q* ≪ *p* in practice. Let *X* and *Y* denote the *n × p* standardized genotype matrix and *n × q* matrix of standardized measured splicing events, respectively. In the first stage of MSG, we use sCCA [24, 69] implemented in the R package “PMA” to identify sparse linear combinations of the columns of *X* and *Y* that are highly correlated with each other. That is, we wish to find vectors *w*_1_ and *u*_1_ that solve the following optimization problem:

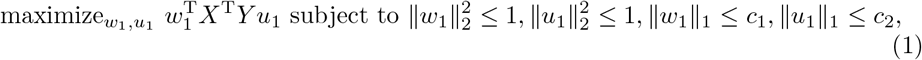

where ||·||_1_ and ||·||_2_ denote the *L*_1_ and *L*_2_ norms, respectively, and *c*_1_ and *c*_2_ are parameters that control the sparsity of *w*_1_ and *u*_1_, respectively. We choose *c*_1_ and *c*_2_ using the default settings of the “CCA” function in “PMA”. Given the selected pair of (*c*_1_, *c*_2_), we obtain subsequent CVs by repeatedly applying the sCCA algorithm (1) to the updated matrix *X*^T^*Y* after regressing out the previous CVs. We repeat this procedure *q* − 1 times to obtain (*w*_2_, *u*_2_), …, (*w*_*q*_, *u*_*q*_). Let *W ≡* (*w*_1_, …, *w*_*q*_) be the *p × q* matrix of SNP weights.

In the second stage of MSG, we test the association between the genetically regulated splicing CVs and the trait of interest using GWAS summary statistics. Specifically, let *z* be the vector of *z*-statistics in the GWAS of trait of interest. The multivariate *z*-statistic for the association between genetically regulated splicing CVs and the trait of interest is *W* ^T^*z*. Under the null hypothesis of no association, it can be shown that *W* ^T^*z* follows a multivariate normal distribution with mean zero and covariance matrix *W* ^T^Σ*W*, where Σ is the *p × p* LD matrix. In practice, we can estimate Σ using an external LD reference panel. A chi-squared test statistic about the gene-trait association can be constructed as

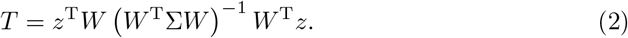

In practice, the splicing events within a gene can be highly correlated, such that the rank of the SNP weight matrix *W* can be less than *q*, and the majority of variations may be explained by a few leading splicing CVs. Consequently, *W* ^T^Σ*W* in expression can be close to singular and its inverse cannot be reliably estimated for many genes. To address this problem, we use the SVD regulation of [19]. Specifically, we compute the pseudo-inverse of *W* ^T^Σ*W* via SVD, decomposing it into its principal components and removing those with small eigenvalues. We use the condition number threshold *λ*_max_/*λ*_*i*_ < 30 to select the number of components, where *λ*_*i*_ and *λ*_*max*_ are the *i*th and maximum eigenvalue of *W* ^T^Σ*W*. Denoting the resulting pseudo-inverse of *W* ^T^Σ*W* as (*W* ^T^Σ*W*)^−^ and substitute it into equation (3), we have

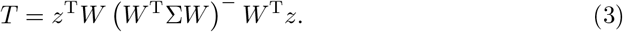

Under the null hypothesis, *T*_2_ follows a 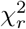 distribution, where *r* is the number of components that contribute to the pseudo-inverse. We test the gene-trait association using a chi-squared test. For each trait and tissue combination, we use Bonferroni correction to determine the genome-wide significance threshold by dividing 0.05 with the number of genes with at least two splicing events in that tissue. This value varies between trait-tissue pairs, and is usually around 0.05*/*10000 = 5 × 10^−6^.

### Simulations

To evaluate the type I error rate and power of the gene-trait association tests, we simulated a training dataset with genetic and splicing data, a GWAS dataset, and a LD reference panel. Then, we conducted gene–trait association tests using our proposed MSG method and the S-MultiXcan, UTMOST, and sCCA+ACAT methods in a variety of realistic scenarios.

To simulate the training dataset with genetic and splicing data, we set *n* = 200, *p* = 300, and *q* = 10. We generated rows of *X* independently from a multivariate normal distribution with mean 0, variance 1, and autoregressive covariance structure determined by *ρ*_*X*_ = 0.1. We generated *Y* from the multivariate linear regression model *Y* = *XB* + *E*, where *B* is a *p × q* matrix of genetic effects on splicing events, and *E* is *n × q* matrix of random errors. Following [70], we factor the effect size matrix *B* into SNP- and splicing event-dependent components, such that *B* = diag(*b*)*D*, where *b* is a *p*-vector of shared genetic effects on all splicing events, diag(*b*) is the *p × p* diagonal matrix expanded by *b*, and *D* is a *p × q* matrix of splicing event-specific effects. We specify the structure of *D* through the following parameters: the number of effect-sharing splicing events (i.e., sharing = 2, 4, 8), the fraction of shared SNPs among non-zero effect SNPs for those effect-sharing splicing events (fixed at 0.3); and the proportion of genetic variants that have non-zero effects on splicing (i.e., sparsity = 1%, 5%, 10%). We generated the elements of *b* independently from a standard normal distribution. We generated the non-zero elements in *D* independently from the uniform distribution on [–1, 1]. We generated the rows of *E* independently from a multivariate normal distribution with mean zero, variance scaled such that the desirable splicing heritability 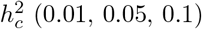 was achieved, and autoregressive covariance structure determined by *ρ*_*E*_ = 0.5.

To simulate the GWAS dataset, we generated a genotype matrix *X*_1_ with 50,000 rows representing the subjects and 300 columns representing the cis-SNPs. We generated the rows of *X*_1_ in a similar manner as we generated *X*. We generated the trait of interest using *Y*_1_ = *X*_1_*Bα* + *ϵ*, where *α* is a *q*-vector of splicing effects on the trait, and *ϵ* is a vector of random errors. Under the null hypothesis of no gene-trait association, we set *α* = 0. Under the alternative hypothesis, we denote the splicing events with non-zero elements in *α* as the “trait-contributing splicing events”, with non-zero values generated independently from a uniform distribution on [−1, 1]. We generated the elements of *E* independently from a normal distribution with mean zero and variance scaled such that the trait heritability was 0.01. We first generated the individual-level dataset and then obtained the GWAS summary statistics. We assumed that only the GWAS summary statistics rather than the individual-level data were available in the subsequent gene-trait association analysis.

We generated an independent genotype matrix *X*_3_ in a similar manner as we generated *X*_1_ and *X*_2_ and used it as an external LD reference panel. We considered two sample sizes for this LD reference panel: 400 (mimicking the 1000 Genomes European reference samples) and 5,000 (mimicking the randomly selected BioVU European samples). Our simulation showed that the MSG method requires more than 400 subjects in the LD reference panel to ensure proper type I error control (Table S1).

We considered a number of realistic scenarios by varying splicing sparsity, splicing heritability, effect-sharing splicing events, and trait-contributing splicing events. When implementing the S-MultiXcan, MSG, and sCCA+ACAT methods, we used their default settings. For type I error evaluation, we used 20,000 replicates for each scenario and used the *p*-value cutoff of 0.05. For power evaluation, we used 2,000 replicates for each scenario and used the *p*-value cutoff of 5 × 10^−6^, which was chosen to mimic the Bonferroni correction in real data applications.

### Compilation of well-known trait-associated gene lists

We obtained AD genes (Supplementary Note Section 1A) from [31]. The authors performed intensive hand-curation to identify confident AD-associated genes from various disease gene resources, including AlzGene, AlzBase, OMIM, DisGenet, DistiLD, UniProt, Open Targets, GWAS Catalog, ROSMAP, and existing literature. We obtained LDL-C genes (Supplementary Note Section 1B) from [43] that included genes from KEGG pathways and existing literature. We obtained a list of genes that influence schizophrenia via splicing (Supplementary Note Section 1C) from [54, 71–73].

### Availability of data and materials

The genotype data for the GTEx project are available on AnVIL (https://anvilproject.org/learn/reference/gtex-v8-free-egress-instructions#downloading-vs-analyzing-in-terra) [74]. Processed GTEx gene expression and splicing data (fully processed, filtered, and normalized splice phenotype matrices in BED format) are downloaded from the GTEx portal (https://gtexportal.org/home/datasets) [75]. With the downloaded GTEx data, we formatted each protein-coding gene with at least two splicing events into a genotype matrix and a splicing events matrix for downstream TWAS analyses. The source of the summary statistics datasets of all GWAS meta-analyses analyzed in this paper can be found in Table S2. The LD matrices for cis-SNPs of each gene from a reference panel of 5,000 randomly selected BioVU samples of European ancestry will be available at the repository Zenodo. The LD reference panel from 1000 Genomes is available at https://data.broadinstitute.org/alkesgroup/FUSION/LDREF.tar.bz2 [76]. In this analysis, we restricted the analysis to SNPs in the HapMap 3 reference panel that are in the LD reference dataset (https://data.broadinstitute.org/alkesgroup/FUSION/LDREF.tar.bz2 [76]) since we are focused on common, well-imputed variants [26]. The code for MSG is available on Github at https://github.com/yingji15/MSG_public [77].

## Supporting information

### AdditionalFile1.pdf

**Supplementary Note 1**. Well-known trait-associated gene lists.

**Fig S1**. AD genes identified via splicing analysis using MSG that would have been missed from expression analysis using S-PrediXcan.

**Fig S2**. LDL-C genes identified via splicing analysis using MSG that would have been missed from expression analysis using S-PrediXcan.

**Fig S3**. Schizophrenia genes identified via splicing analysis using MSG that would have been missed from expression analysis using S-PrediXcan.

**Table S1**. Comparison of type I error for MSG using individual GWAS, MSG with GWAS summary statistics and reference genome of 400 and 5000 individuals in simulation.

### AdditionalFile2.xlsx

**Tables S2-S16**. Summary of MSG application to 14 human traits. The source of GWAS and MSG identified trait-associated genes are provided.

**Tables S17-S19**. MSG identified genes that are within 500 kb distance to GWAS significant loci in AD, LDL-C and schizophrenia.

**Tables S20-S22**. MSG identified genes that overlap with S-PrediXcan identified genes in AD, LDL-C, and schizophrenia.

## Acknowledgments

We thank Drs. Nancy Cox, Yi Jiang, and Dan Zhou for their helpful discussions. This work was conducted in part using the resources of the Advanced Computing Center for Research and Education at Vanderbilt University, Nashville, TN.

